# Mesophotic gorgonian corals evolve multiple times and faster than deep and shallow lineages

**DOI:** 10.1101/2020.12.17.422867

**Authors:** Juan A. Sánchez, Fanny L. González-Zapata, Carlos Prada, Luisa F. Dueñas

## Abstract

Mesophotic Coral Ecosystems (MCEs) develop on a unique environment, where abrupt environmental changes take place. Using a time-calibrated molecular phylogeny (mtDNA: mtMutS), we examined the lineage membership of mesophotic gorgonian corals (Octocorallia: Cnidaria) in comparison to shallow and deep-sea lineages of the wider Caribbean-Gulf of Mexico and the Tropical Eastern Pacific. Our results show mesophotic gorgonians originating multiple times from old deep-sea octocoral lineages, whereas shallow-water species comprise younger lineages. The mesophotic gorgonian fauna in the studied areas are related to their zooxanthellate shallow-water counterparts in only two clades (Gorgoniidae and Plexauridae), where the shallow-deep gradient could serve as a driver of diversification. Interestingly, mesophotic clades have diversified faster than either shallow or deep clades. One of this groups with fast diversification is the family Ellisellidae, a major component of the mesophotic gorgonian coral assemblage worldwide.

## 2. Introduction

Gorgonian corals (Cnidaria: Octocorallia) generate a unique seascape in the shallow-water communities of the Western Atlantic and adjacent seas, including Mesophotic Coral Communities (MCEs). Their tall branching colonies, sometimes reaching high densities, form an animal forest of great diversity from shallow to mesophotic and deeper ecosystems [1,2]. Dense mesophotic gorgonian assemblages thrives at both coasts of tropical and sub-tropical America [3], the IndoPacific [2], Brazil [4], the west coast of Africa [5], and even some temperate areas in the Atlantic [6,7] and the Mediterranean Sea [8]. MCEs develop on an exceptional environment limiting the growth of many coral species and leading to many depth-specialist adaptation [9]. MCEs include a diverse gorgonian fauna, yet it is unknown if this fauna is the extension of shallow or deep-sea communities or whether mesophotic octocorals comprised different evolutionary lineages [2]; moreover, the role of this ecosystem on octocoral diversification is unknown.

In this study, we reconstructed a time-calibrated molecular phylogeny for 242 gorgonian coral species using mtDNA (mtMutS), including numerous new sequences of mesophotic gorgonian corals from the Caribbean Sea (down to ∼120 m) and many valuable sequences in Genbank from related species in the Gulf of Mexico [10], eastern tropical Pacific [11–13], western Atlantic [14], and Indo-Pacific [15]. Collectively, this mtMutS database comprise a comprehensive set of sampling otherwise impossible to acquire for a single study. We tested whether mesophotic lineages descent from a single ancestor from either shallow or deep-water areas or if instead have multiple origins. We also compared rates of diversification across shallow, mesophotic and deep-water gorgonians.

## 3. Materials and Methods

Using Closed-Circuit Rebreather-CCR and Trimix, we surveyed gorgonian corals from 115 m up to 45 m in MCEs in three locations in the Colombian Caribbean: San Andrés Island (Archipelago of San Andrés, Providencia and Santa Catalina), Barú island diapiric banks and the Deep-sea Corals National Park (both near Cartagena). A dry voucher for each colony is available at the Museo de Historia Natural ANDES (ANDES-IM 4132 to ANDES-IM 4802). Research and collection of specimens were approved by the National Environmental Licensing Authority (ANLA, Spanish acronym): Collection Framework Agreement granted to Universidad de los Andes through resolution 1177 of October 9, 2014 - IBD 0359. Since we did the collections during previous studies, detailed information on the sites and study areas is already available **[16–18]**. Together with our new material, we examined their phylogenetic affiliations in comparison to shallow and deep-sea lineages of the wider Caribbean-Gulf of Mexico and Tropical Eastern Pacific using the available information (See supplementary table 1).

Samples were fixed in both Ethanol 95% and DMSO. Total genomic DNA of each specimen was extracted from about 5 mm^2^ of tissue following a standard CTAB Phenol:Chloroform:Isoamyl Alcohol protocol [19]. DNA quality was assessed in 1% agarose gel electrophoresis in 1X TBE buffer. Gels were dyed with ethidium bromide and visualized in a Gel Doc^™^ XR (Biorad, U.S.). An approximate estimation of concentration in ng •l^-1^ and purity (260/280 and 260/230 ratios) of each DNA sample was assessed with a NanoDrop (Thermo Scientific, U.S.). The mtMutS region was targeted using the protocols described in the literature [20].

Phylogenetic relationships and times of divergence between shallow and deep-sea gorgonian lineages were co-estimated using BEAST ver. 1.8.2. Divergence times were estimated using a relaxed molecular clock with log-normal uncorrelated rates and assuming a Birth-Death Incomplete Sampling speciation tree prior. The analysis was run four independent times under a GTR model and used 10^7^ generations and default heating values on three Metropolis-coupled chains. Trees and parameters were sampled every 1,000 generations and the first 10% of the samples were discarded as burn-in. We used Tracer ver. 1.8 to check for adequate convergence and confirmed effective sample size (ESS) greater than 200. LogCombiner ver. 1.8 and TreeAnnotator ver. 1.8 were used to combine and summarize tree files, obtain a maximum clade credibility consensus tree, and calculate 95% credibility intervals. We also ran the analysis on an empty dataset, sampling from the prior distribution to evaluate the influence of the priors on the posterior distribution estimates [21]. We followed a time-calibrated molecular approach using fossil calibration points [20]. As calibration points, we employed the oldest known fossil for the families Elliseliidae (Kuzmicheva 1987) and Keratoisidinae [22], and for the genus *Eunicella* [23]. The minimum age of each fossil was treated as a minimum constraint on the age of the stem group node using a log-normal distributed prior. The standard deviation was calculated in such a way that 95% of the probability density lies between the minimum constraint and the oldest date of the geological range of the fossil. Letters correspond to monophyletic lineages explained in the text.

To test if clades from different depths (shallow, mesophotic of deep) differ in their diversification rates, we used the multi-state character extension (*MuSSE*) model as implemented in the package *diversitree* [24]. Initially we fit a “null” model, in which all birth and death rates were equal between states, the character evolution was ordered (shallow <-> mesophotic <-> deep), and there is a single character transition rate. We then fitted models in which only the speciation rate (•) varied between states, only the extinction rate (•) varied, and finally, one in which neither • nor • vary, but the transition rates differ between types of transitions. We then fitted a more complex model in which all rates of speciation and extinction depended on the character state for our multi-state character. To rank and choose among the different models with speciation and extinction rates, we used the Akaike information criterion (AIC). Using information theory and AIC, we computed the relative weight of evidence in favor of each of our different hypotheses using AIC weights and chose the best model [25]. With the best (selected) model, we run a Bayesian MCMC. We run our chain with 9,000 steps.

In addition, and to understand the evolution of habitat use among these gorgonians, we estimated habitat use values for ancestral nodes in the inferred phylogenetic tree. We modeled our characters using a discrete approach using a continuous-time Markov chain model commonly known as the Mk model. We then fitted a single-rate model and reconstructed ancestral states at internal nodes in the phylogeny. We used the function *lik*.*anc* to estimate the marginal ancestral states. As an alternative way to reconstruct states at ancestral nodes, we sampled character histories from their posterior probability distribution using an MCMC approach, known as the “stochastic character mapping” [26] with the *make*.*simmap* function in package “*phytools*” [27]. In this latter approach, we obtained a sample of histories for our discrete character’s evolution on the phylogeny - rather than a probability distribution for the character at nodes. Since a single reconstruction is meaningless, we iterated the process 1,000 times and evaluated the distribution from these stochastic maps. To generate a summary of these maps, we estimated the number of changes of each type, the proportion of time spent in each state, and the posterior probabilities that each internal node is in each state, under our model.

## 4. Results

The obtained time-calibrated phylogeny showed high support values for all studied lineages at the genus level and major recognized clades (See supplementary Fig. 1). Overall, there were trends separating shallow, mesophotic, and deep gorgonian species but multiple shifts to different depth ranges occurred. Colonization of Caribbean MCEs happened even at the oldest octocoral lineages, i.e., stem age >∼100 MYA, such as *Trychogorgia lyra* (Chrysogorgiidae) a species within the deep-sea clade of highly calcified octocorals (Calcaxonia) (Fig. 1, clade A). Despite gorgonian corals forming similar branching tree-like colonies and habitat-forming characteristics, they are a polyphyletic group including old deep-sea lineages lacking hard or proteinaceous skeletons, also known as scleraxonians (Fig. 1, clades B and C) closely related with soft corals, which include common Caribbean MCE members such as *Iciligorgia* and *Diodogorgia* [28,29]. Preceded by a clade of deep-sea stoloniferous octocorals, the true gorgonian corals (Fig. 1, clade D), i.e., with an axial proteinaceous skeleton, grouped in a large younger clade (< 100 MYA stem age), contained modest phylogenetic resolution with several patterns that we describe below.

**Figure 1.**
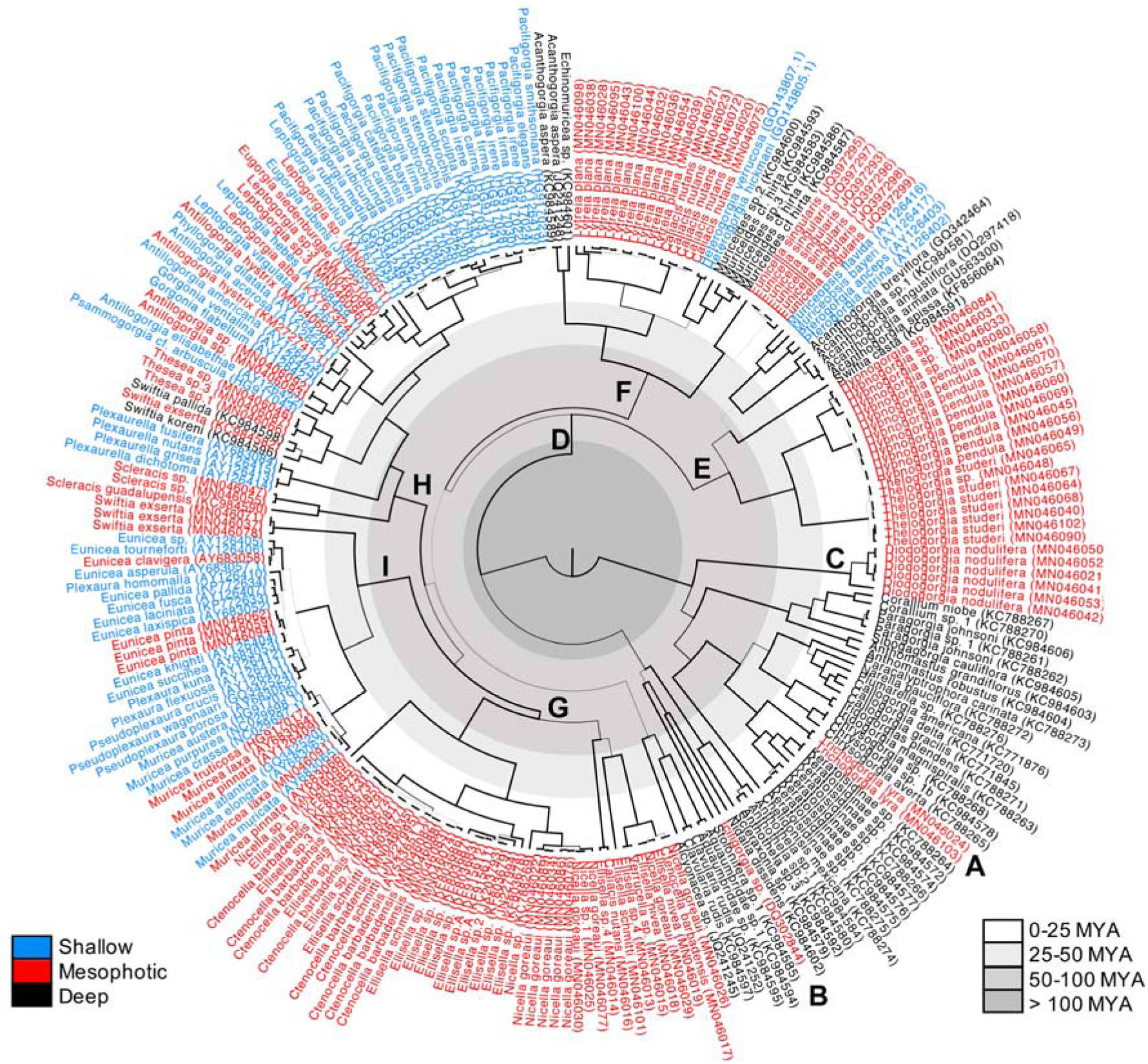
Time-calibrated phylogeny from 242 gorgonian coral species from shallow-, mesophotic-, and deep-water gorgonian corals from the Caribbean-Gulf of Mexico and the eastern tropical Pacific (mtMutS). Branch line width represents posterior probability support; thicker lines are supports >0.9. Important clades are labeled A-I correspond, which correspond to monophyletic lineages explained in the text. (See supplementary Figure 1 for the chronogram with error bars)

First, we find a clade including Acanthogorgiidae (*Acanthogorgia* spp.), *Hypnogorgia* and the family Keroeididae (*Thelogorgia*) (Fig. 1, clade E), along with two shallow-water groups, *Muriceopsis* and *Pterogorgia*, that are usually attracted to disparate clades in the octocoral phylogeny probably due to long branch attraction [28,30]. Second, part of the paraphyletic family Plexauridae in the clade known as ‘Stenogorgiinae’ [31] arises in the tree, most reaching mesophotic depths, but also found at depths below 200 m such as *Lytreia, Muriceides, Heterogorgia* (only shallow-water), *Caliacis, Echinomuricea* and *Eunicella singularis* (Fig. 1, clade F). Third, we notice Ellisellidae clade, a major component of the mesophotic gorgonian coral assemblage, attaining high densities in the upper MCE range [2,18,32], and the only group where MCEs promoted its entire diversification (Figure 1, clade G). This group, extending also to deeper ecosystems, is the only one found in MCEs worldwide [33], and its simultaneous parallel evolution [15] suggests that MCEs could be an important factor in their diversification.

Last, we see the shallow-water gorgonian corals, including all zooxanthellate species from the Caribbean, and the azooxanthellate, including aposymbiotic *Muricea* [33], from the Tropical Eastern Pacific, appear in two clades that we can assign to the families Plexauridae (in part) and Gorgoniidae, major components of the shallow-water communities (Fig. 1, clades H and I). Plexauridae includes mesophotic-associated genera such as *Scleracis, Swiftia* (in part), and *Thesea*, some groups including shallow and mesophotic groups like *Leptogorgia* and *Eugorgia* [34], azooxanthellate shallow-water *Pacifigorgia* and *Psammogorgia*, the zooxanthellate *Plexaurella, Gorgonia, Phyllogorgia* and *Antillogorgia*, the latter includes some mesophotic gorgonian corals (Fig. 1, clade H). The Plexauridae clade has the Caribbean *Swiftia exserta* as sister clade, with species in both Caribbean and the Tropical Eastern Pacific, and diverse Caribbean zooxanthellate shallow-water groups, *Pseudoplexaura, Plexaura, Muricea* and *Eunicea* (Fig. 1, clade I), which includes mostly mesophotic species [2]. Some of these symbiotic mesophotic species (e.g., *Muricea laxa* and *Antillogorgia hystrix*), are likely the product of shallow-deep ecological divergence, similar to the incipient cases of *Eunicea flexuosa* and *Antillogorgia bipinnata* [35,36], but today reaching depths below 40 m. There were multiple unresolved relationships in the octocoral phylogeny observed recurrently, even with the largest amount of phylogenetic information [37], which together with the placement of Ellisellidae, are beyond the scope of this article and deserve further systematic revision [29].

Overall speciation was faster in mesophotic and shallow-water gorgonian clades. Yet, less extinction was detected in the deep-sea lineages. Remarkable, net diversification rates were faster in mesophotic lineages followed by shallow and deep clades (Fig. 2). In addition, mesophotic gorgonian corals had multiple deep-sea origins. Shallow-water gorgonian lineages, which are more abundant in the sampling and apparently more speciose, were restricted to less clades than mesophotic gorgonians, which revolutionize from several deep-sea ancestors (Figs. 1-2).

**Figure 2.**
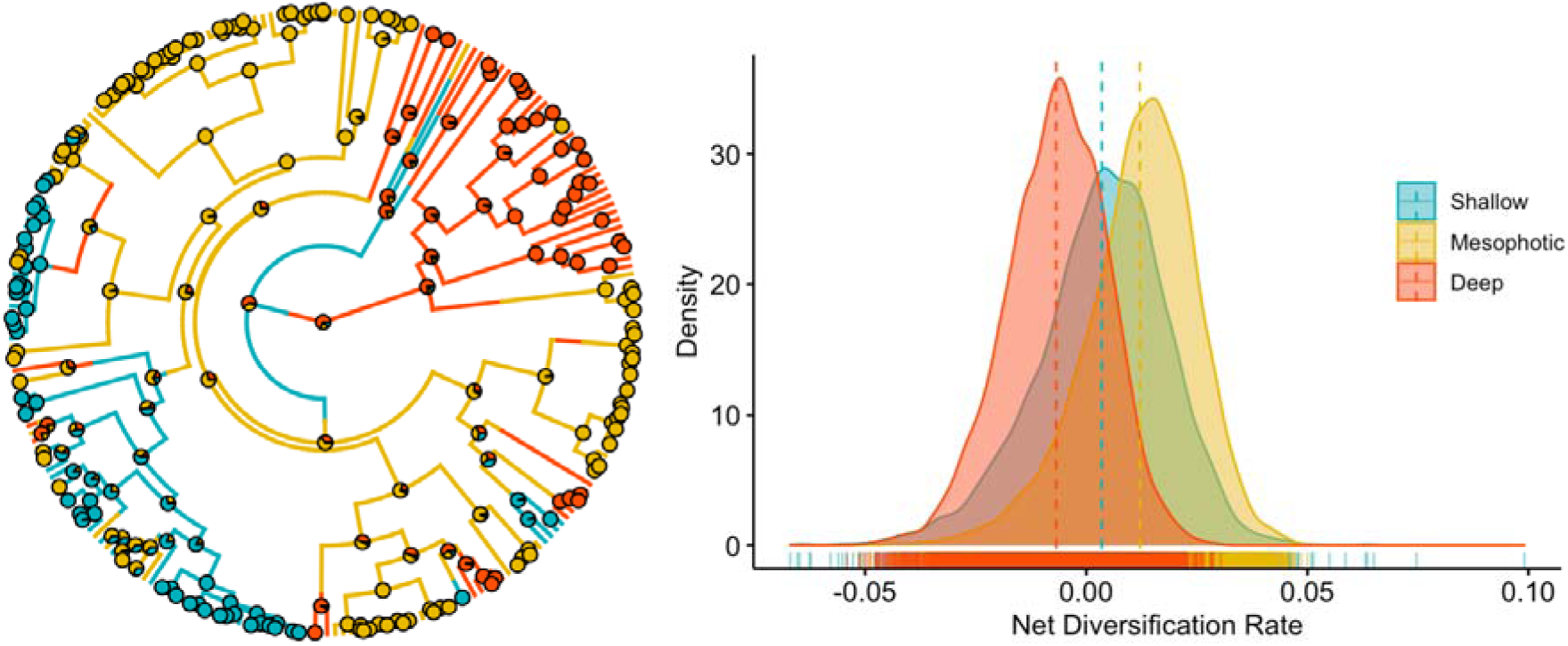
A. The phylogeny with a discreate character map based on a summary of 1,000 stochastic maps generated from modeling of the evolution of habitat use. Branches are colored depending on the habitat utilization by the different gorgonians. B. Rates of net diversification for shallow, mesophotic and deep-water gorgonians. Probability density plots are based on 9,000 MCMC samples of the full MuSSE model.

## 5. Discussion

The little genetic divergence within species from shallow-water genera in the Caribbean and Tropical Eastern Pacific have been noted by several authors **[12**,**30**,**31**,**38–41]**. Apart from the families Plexauridae and Gorgoniidae, where ecological divergence is suspected as the gorgonian assemblage colonizes the upper MCEs (30-60 m), the relationship between shallow-mesophotic species was rarely observed in closely related species in most clades. This pattern is also found in reef fishes and scleractinian corals [42]. Although reef-building scleractinian corals can colonize deeper into MCEs [16,43,44], the azooxathellate cup corals replaces the hermatypic coral assemblage [18] and belongs to very old (>77 MYA) Scleractinian lineages, dominating this environment and even deeper waters [45]. Our study also supports the notion that gorgonian corals (“branching holaxonians”) had the fast evolutionary rates among all anthozoans [46].

The families Plexauridae and Gorgoniidae showed replicated patterns of sister species segregated by shallow and mesophotic habitats consistent with recent research showing how depth plays a major role in the diversification of reef organisms [36,47]. The shallow-deep gradient creates an intermediate scenario between adaptive plasticity and local adaptation, which is common several Caribbean species [3,35]. Estimates of young diversification between pairs of habitat-segregated pairs of species is consistent with recent demographic models inferred from genomic data [48]. Our phylogenetic reconstruction suggests that such a shallow-mesophotic diversification has occurred at least nine times in these two families, pointing out by the first time, to the major role and the macroevolutionary magnitude of depth/light promoting the formation of new species in the Caribbean. We suspect that ecological specialization mediated by immigrant inviability, as suggested previously [36], mediates the formation of these young pairs of segregated species in shallow and mesophotic habitats.

Mesophotic gorgonian corals in the Caribbean, also excluding Plexauridae and Gorgoniidae, have close memberships with deep-sea groups and can be located at the shallower records of those lineages [1,49]. Gorgonians living at mesophotic depths (45-182 m) exceed the geographical/latitudinal bounds of shallow-water species [50], which supports the idea of their independent evolutionary history. Two families, which most species distribute within the MCE range, Keroeididae, with all species of the genus *Thelogorgia* [51] and Ellisellidae, which suggest the MCE depth range and environment can be considered an important feature for octocoral diversification. Previous observations in the upper mesophotic zone (30-60m) from Caribbean reefs suggested that younger gorgonian species lineages are replaced by older lineages characterized by phylogenetically dispersed species, which have thinner branches and smaller polyps than shallow-water species [14]. Likewise, polyp density decreases with depth in gorgonian corals [52], which has been hypothesized as a response to an increasing microbial metabolism due low water-motion and anoxia with depth [2]. In general, mesophotic gorgonian corals in the Caribbean are not related to their shallow-water counterparts and consequently, mesophotic depths do not serve as a refuge for shallow gorgonians excluding Plexauridae and Gorgoniidae, where MCEs are being colonized back. Interestingly, mesophotic clades seem to have faster diversification rates than both shallow and deep-water gorgonians (Fig. 2).

## Acknowledgments

The support from Bluelife dive shop (family Garcia) was fundamental to accomplish this study. The San Andres Hospital kindly supplied medical oxygen for CCR. We are very grateful with Gregg Stanton, Wakulla Dive Center, for continuing support and advise for deep diving. We are thankful with Nacor Bolaños, Julio Andrade, Fabian García, Santiago Herrera, Mariana Gnecco, Manu Forero, Deibis Seguro, Oscar Ruiz, Federico Botero and Camilo Martinez for fieldwork support. We recognize the participation and support from local communities.

## Ethical Statement

Our project grant (COLCIENCIAS No. 120465944147) was reviewed by Universidad de los Andes ethical committee and found it “low risk”. All permits and collecting protocols met the Colombian laws.

## Data Accessibility

DNA sequences: **New sequences have GenBank Accession Numbers MN046013-MN046103**.

## Authors’ Contributions

JAS, FLG-Z and LFD conceived the study. JAS, FLG-Z, CP and LFD ran the analyses. JAS wrote the manuscript with the help of FLG-Z, LFD and CP. All authors approved the final version of the article.

