## supplementary table 1S for "Mesophotic gorgonian corals evolve multiple times and faster than deep and shallow lineages"

**Supplementary Table 1S**. Metadata from sequences used in the study.

| GenBank ACC No. | Type | Depth | Species | Specimen_voucher | Country |
|---|---|---|---|---|---|
| *New sequences generated for this study (D=deep-water; M=mesophotic; S=shallow-water)* | | | | | |
| MN046063 | M | 60m | Antillogorgia hystrix | ANDES-IM 4262 | Colombia |
| MN046092 | M | 45m | *Antillogorgia* sp. | ANDES-IM 4459 | Colombia |
| MN046093 | M | 45m | *Antillogorgia* sp. | ANDES-IM 4461 | Colombia |
| MN046016 | M | 37m | *Caliacis nutans* | ANDES-IM 4661 | Colombia |
| MN046020 | M | 35m | *Caliacis nutans* | ANDES-IM 4658 | Colombia |
| MN046023 | M | 80m | *Caliacis nutans* | ANDES-IM 4347 | Colombia |
| MN046027 | M | 80m | *Caliacis nutans* | ANDES-IM 4348 | Colombia |
| MN046072 | M | 95m | *Caliacis nutans* | ANDES-IM 4801 | Colombia |
| MN046075 | M | 95m | *Caliacis nutans* | ANDES-IM 4645 | Colombia |
| MN046017 | M | 37m | *Ctenocella barbadensis* | ANDES-IM 4146 | Colombia |
| MN046035 | M | 80m | *Ctenocella barbadensis* | ANDES-IM 4132 | Colombia |
| MN046082 | M | 90m | *Ctenocella barbadensis* | ANDES-IM 4139 | Colombia |
| MN046085 | M | 85m | *Ctenocella barbadensis* | ANDES-IM 4500 | Colombia |
| MN046088 | M | 85m | *Ctenocella barbadensis* | ANDES-IM 4143 | Colombia |
| MN046021 | M | 45m | *Diodogorgia nodulifera* | ANDES-IM 4368 | Colombia |
| MN046041 | M | 72m | Diodogorgia nodulifera | ANDES-IM 4795 | Colombia |
| MN046042 | M | 72m | Diodogorgia nodulifera | ANDES-IM 4796 | Colombia |
| MN046050 | M | 72m | *Diodogorgia nodulifera* | ANDES-IM 4366 | Colombia |
| MN046052 | M | 72m | *Diodogorgia nodulifera* | ANDES-IM 4367 | Colombia |
| MN046053 | M | 72m | *Diodogorgia nodulifera* | ANDES-IM 4797 | Colombia |
| MN046018 | M | 37m | *Ellisella nivea* | ANDES-IM 4363 | Colombia |
| MN046019 | M | 37m | *Ellisella nivea* | ANDES-IM 4365 | Colombia |
| MN046083 | M | 90m | *Ellisella schmitti* | ANDES-IM 4440 | Colombia |
| MN046101 | M | 50m | *Ellisella schmitti* | ANDES-IM 4439 | Colombia |
| MN046074 | M | 95m | *Ellisella* sp. | ANDES-IM 4497 | Colombia |
| MN046073 | M | 95m | *Ellisella* sp3 | ANDES-IM 4800 | Colombia |
| MN046013 | M | 60m | *Ellisella* sp4 | ANDES-IM 4494 | Colombia |
| MN046014 | M | 60m | *Ellisella* sp4 | ANDES-IM 4495 | Colombia |
| MN046079 | M | 115m | *Ellisella* sp7 | ANDES-IM 4498 | Colombia |
| MN046081 | M | 115m | *Ellisella* sp7 | ANDES-IM 4499 | Colombia |
| MN046059 | M | 80m | *Eunicea pinta* | ANDES-IM 4409 | Colombia |
| MN046062 | M | 60m | *Eunicea pinta* | ANDES-IM 4410 | Colombia |
| MN046094 | M | 45m | *Eunicea pinta* | ANDES-IM 4406 | Colombia |
| MN046045 | M | 72m | *Hypnogorgia pendula* | ANDES-IM 4620 | Colombia |
| MN046049 | M | 72m | *Hypnogorgia pendula* | ANDES-IM 4624 | Colombia |
| MN046056 | M | 80m | *Hypnogorgia pendula* | ANDES-IM 4369 | Colombia |
| MN046057 | M | 80m | *Hypnogorgia pendula* | ANDES-IM 4639 | Colombia |
| MN046058 | M | 80m | *Hypnogorgia pendula* | ANDES-IM 4640 | Colombia |
| MN046060 | M | 80m | *Hypnogorgia pendula* | ANDES-IM 4641 | Colombia |
| MN046061 | M | 80m | *Hypnogorgia pendula* | ANDES-IM 4370 | Colombia |
| MN046069 | M | 95m | *Hypnogorgia pendula* | ANDES-IM 4371 | Colombia |

| | | | | | |
|---|---|---|---|---|---|
| MN046070 | M | 95m | *Hypnogorgia pendula* | ANDES-IM 4644 | Colombia |
| MN046031 | M | 80m | *Hypnogorgia* sp. | ANDES-IM 4633 | Colombia |
| MN046033 | M | 80m | *Hypnogorgia* sp. | ANDES-IM 4634 | Colombia |
| MN046080 | M | 115m | *Hypnogorgia* sp. | ANDES-IM 4372 | Colombia |
| MN046084 | M | 85m | *Hypnogorgia* sp. | ANDES-IM 4646 | Colombia |
| MN046051 | M | 72m | *Leptogoragia* sp. | ANDES-IM 4504 | Colombia |
| MN046096 | M | 50m | *Leptogorgia* sp3 | ANDES-IM 4505 | Colombia |
| MN046097 | M | 50m | *Leptogorgia* sp3 | ANDES-IM 4506 | Colombia |
| MN046028 | M | 80m | *Lytreia plana* | ANDES-IM 4421 | Colombia |
| MN046032 | M | 80m | *Lytreia plana* | ANDES-IM 4422 | Colombia |
| MN046036 | M | 77m | *Lytreia plana* | ANDES-IM 4629 | Colombia |
| MN046038 | M | 77m | *Lytreia plana* | ANDES-IM 4418 | Colombia |
| MN046039 | M | 77m | *Lytreia plana* | ANDES-IM 4476 | Colombia |
| MN046043 | M | 72m | *Lytreia plana* | ANDES-IM 4605 | Colombia |
| MN046044 | M | 66m | *Lytreia plana* | ANDES-IM 4513 | Colombia |
| MN046054 | M | 72m | *Lytreia plana* | ANDES-IM 4475 | Colombia |
| MN046095 | M | 85m | *Lytreia plana* | ANDES-IM 4802 | Colombia |
| MN046098 | M | 50m | *Lytreia plana* | ANDES-IM 4511 | Colombia |
| MN046100 | M | 50m | *Lytreia* plana | ANDES-IM 4512 | Colombia |
| MN046091 | M | 45m | *Muricea laxa* | ANDES-IM 4794 | Colombia |
| MN046022 | M | 80m | *Nicella goreaui* | ANDES-IM 4212 | Colombia |
| MN046026 | M | 80m | *Nicella goreaui* | ANDES-IM 4213 | Colombia |
| MN046029 | M | 80m | *Nicella goreaui* | ANDES-IM 4522 | Colombia |
| MN046030 | M | 80m | *Nicella goreaui* | ANDES-IM 4214 | Colombia |
| MN046034 | M | 80m | *Nicella goreaui* | ANDES-IM 4215 | Colombia |
| MN046076 | M | 115m | *Nicella goreaui* | ANDES-IM 4655 | Colombia |
| MN046077 | M | 115m | *Nicella goreaui* | ANDES-IM 4219 | Colombia |
| MN046086 | M | 85m | *Nicella goreaui* | ANDES-IM 4220 | Colombia |
| MN046089 | M | 85m | *Nicella goreaui* | ANDES-IM 4221 | Colombia |
| MN046025 | M | 80m | *Nicella* sp1 | ANDES-IM 4521 | Colombia |
| MN046099 | M | 50m | *Nicella* sp1 | ANDES-IM 4519 | Colombia |
| MN046055 | M | 87m | *Scleracis* sp. | ANDES-IM 4626 | Colombia |
| MN046047 | M | 72m | *Scleracis* sp. | ANDES-IM 4622 | Colombia |
| MN046037 | M | 77m | *Swiftia exserta* | ANDES-IM 4609 | Colombia |
| MN046071 | M | 95m | *Swiftia exserta* | ANDES-IM 4193 | Colombia |
| MN046078 | M | 115m | *Swiftia exserta* | ANDES-IM 4194 | Colombia |
| MN046040 | M | 72m | *Thelogorgia studeri* | ANDES-IM 4615 | Colombia |
| MN046064 | M | 67m | *Thelogorgia studeri* | ANDES-IM 4665 | Colombia |
| MN046065 | M | 95m | *Thelogorgia studeri* | ANDES-IM 4666 | Colombia |
| MN046067 | M | 95m | *Thelogorgia studeri* | ANDES-IM 4667 | Colombia |
| MN046068 | M | 95m | *Thelogorgia studeri* | ANDES-IM 4668 | Colombia |
| MN046090 | M | 85m | *Thelogorgia studeri* | ANDES-IM 4670 | Colombia |
| MN046102 | M | 50m | *Thelogorgia studeri* | ANDES-IM 4664 | Colombia |
| MN046046 | M | 72m | *Thesea* sp. | ANDES-IM 4621 | Colombia |
| MN046066 | M | 95m | *Thesea* sp1 | ANDES-IM 4643 | Colombia |

| | | | | | |
|---|---|---|---|---|---|
| MN046087 | M | 85m | *Thesea* sp3 | ANDES-IM 4647 | Colombia |
| MN046024 | M | 80m | *Trichogorgia lyra* | ANDES-IM 4342 | Colombia |
| MN046103 | M | 50m | *Trichogorgia lyra* | ANDES-IM 4653 | Colombia |
| MN046048 | M | 72m | Unknown octocoral | ANDES-IM 4617 | Colombia |
| MN046015 | M | 60m | *Verrucella* sp. | ANDES-IM 4654 | Colombia |
| *Sequences obtained from GenBank* | | | | | |
| DQ297418 | D | | *Acanthogorgia angustiflora* | | |
| GU563300 | D | | *Acanthogorgia armata* | | |
| JQ241248 | D | | *Acanthogorgia aspera* | | |
| KC984589 | D | | *Acanthogorgia aspera* | | |
| GQ342464 | D | | *Acanthogorgia breviflora* | | |
| KC984581 | D | | *Acanthogorgia* sp1 | | |
| KF856064 | D | | *Acanthogorgia spissa* | | |
| JQ241245 | D | | *Alcyonacea* sp. | | |
| KC984603 | D | | *Anthomastus grandiflorus* | | |
| KC984604 | D | | *Anthomastus robustus* | | |
| KC984584 | D | | *Anthothela* sp1 | | |
| KC984580 | D | | *Anthothela* sp2 | | |
| KC984592 | D | | *Anthothela* sp3 | | |
| KM272741.1 | S | | *Antillogorgia hystrix* | | |
| KC984595 | D | | *Aquaumbridae* sp. | | |
| KC984594 | D | | *Aquaumbridae* sp1 | | |
| KC771876 | D | | *Callogorgia americana* | | |
| KC771720 | D | | *Callogorgia delta* | | |
| KC771845 | D | | *Callogorgia gracilis* | | |
| KC788274 | D | | *Chelidonisis mexicana* | | |
| KC788265 | D | | *Chrysogorgia averta* | | |
| KC788268 | D | | *Chrysogorgia* sp. | | |
| KC984578 | D | | *Chrysogorgia* sp1b | | |
| JQ241252 | D | | *Clavularia rudis* | | |
| KC984597 | D | | *Clavularia rudis* | | |
| KC788267 | D | | *Corallium niobe* | | |
| KC788270 | D | | *Corallium* sp1 | | |
| AY533651.1 | M | | *Ctenocella barbadensis* | | |
| KF803673 | M | | *Ctenocella barbadensis* | | |
| KF803674 | M | | *Ctenocella barbadensis* | | |
| KF803675 | M | | *Ctenocella barbadensis* | | |
| JN227995 | M | | *Ctenocella schmitti* | | |
| KC984601 | D | | *Echinomuricea* sp. | | |
| KF803677 | M | | *Ellisella schmitti* | | |
| JN227994 | M | | *Ellisella* sp. | | |
| KF803678 | M | | *Ellisella* sp. | | |
| KF803679 | M | | *Ellisella* sp. | | |
| KF803683 | M | | *Ellisella* sp. | | |
| KF803685 | M | | *Ellisella* sp. | | |

| | | |
|---|---|---|
| KF803686 | M | *Ellisella* sp. |
| KF803687 | M | *Ellisella* sp. |
| JX203793 | M | *Ellisella* sp2 |
| KF915611 | M | *Ellisella* spA |
| KF915612 | M | *Ellisella* spA |
| JN866557.1 | M | *Eugorgia rubens* |
| LT221096 | M | *Eugorgia siedenburgae* |
| AY683057.1 | S | *Eunicea asperula* |
| AY683058 | S | *Eunicea clavigera* |
| AY126407 | S | *Eunicea fusca* |
| AY126404 | S | *Eunicea knighti* |
| KP772633 | S | *Eunicea laciniata* |
| AY683052 | S | *Eunicea laxispica* |
| KP772634 | S | *Eunicea pallida* |
| AY126405 | S | *Eunicea* sp. |
| AY683053 | S | *Eunicea succinea* |
| AY126406 | S | *Eunicea tourneforti* |
| JQ397293 | M | *Eunicella singularis* |
| JQ397295 | M | *Eunicella singularis* |
| JQ397296 | M | *Eunicella singularis* |
| JQ397297 | M | *Eunicella singularis* |
| JQ397298 | M | *Eunicella singularis* |
| JQ397299 | M | *Eunicella singularis* |
| AY126427 | S | *Gorgonia flabellum* |
| AY126425 | S | *Gorgonia ventalina* |
| GQ143805.1 | S | *Heterogorgia hickmani* |
| GQ143807.1 | S | *Heterogorgia verrucosa* |
| DQ302844 | S | *Iciligorgia* sp. |
| KC788263 | D | *Iridogorgia magnispiralis* |
| KC788271 | D | *Iridogorgia splendens* |
| KC788264 | D | *Keratoisidinae* sp. |
| KC788266 | D | *Keratoisidinae* sp. |
| KC788275 | D | *Keratoisidinae* sp. |
| KC984572 | D | *Keratoisidinae* sp. |
| KC984574 | D | *Keratoisidinae* sp. |
| KC984575 | D | *Keratoisidinae* sp. |
| KC984576 | D | *Keratoisidinae* sp. |
| KC984577 | D | *Keratoisidinae* sp. |
| KX767324 | M | *Leptogorgia alba* |
| AY268459.1 | S | *Leptogorgia hebes* |
| KX767322.1 | S | *Leptogorgia ramulus* |
| AY126418 | S | *Leptogorgia virgulata* |
| GQ342520 | S | *Muricea atlantica* |
| HG917016 | S | *Muricea* cf. *austera* |
| NC029697.1 | S | *Muricea crassa* |

| | | |
|---|---|---|
| AY683063 | S | *Muricea elongata* |
| HG917017.1 | S | *Muricea fruticosa* |
| AY683064 | S | *Muricea laxa* |
| AY683065 | S | *Muricea muricata* |
| AY126409 | S | *Muricea pinnata* |
| AY683066 | M | *Muricea pinnata* |
| NC029698.1 | S | *Muricea purpurea* |
| KC984586 | D | *Muriceides* cf. *hirta* |
| KC984587 | D | *Muriceides* cf. *hirta* |
| KC984593 | D | *Muriceides* cf. *hirta* |
| KC984600 | D | *Muriceides* sp2 |
| KC984583 | D | *Muriceides* sp3 |
| AY126417 | S | *Muriceopsis bayeri* |
| AY126416 | S | *Muriceopsis flavida* |
| KC788272 | D | *Narella pauciflora* |
| KC788269 | M | *Nicella* sp. |
| KC984602 | D | *Nidalia dissidens* |
| HG917044 | S | *Pacifigorgia bayeri* |
| HG917021 | S | *Pacifigorgia cairnsi* |
| HG917041 | S | *Pacifigorgia cairnsi* |
| HG917020 | S | *Pacifigorgia catedralensis* |
| AY126419 | S | *Pacifigorgia elegans* |
| HG917022 | S | *Pacifigorgia firma* |
| HG917028 | S | *Pacifigorgia firma* |
| HG917029 | S | *Pacifigorgia firma* |
| HG917024 | S | *Pacifigorgia irene* |
| HG917030 | S | *Pacifigorgia irene* |
| HG917031.1 | S | *Pacifigorgia irene* |
| GQ342497 | S | *Pacifigorgia media* |
| HG917027 | S | *Pacifigorgia rubicunda* |
| HG917032 | S | *Pacifigorgia rubicunda* |
| HG917033.1 | S | *Pacifigorgia rubicunda* |
| KX767343.1 | S | *Pacifigorgia sculpta* |
| HG917023 | S | *Pacifigorgia smithsoniana* |
| AY126420 | S | *Pacifigorgia stenobrochis* |
| HG917018 | S | *Pacifigorgia stenobrochis* |
| HG917026 | S | *Pacifigorgia stenobrochis* |
| KC788273 | D | *Paracalyptrophora carinata* |
| KC788262 | D | *Paragorgia johnsoni* |
| KC984606 | D | *Paragorgia johnsoni* |
| KC788261 | D | *Paragorgia* sp1 |
| AY126428.1 | S | *Phyllogorgia dilatata* |
| AY126411 | S | *Plexaura flexuosa* |
| AY126410 | S | *Plexaura homomalla* |
| AY126412 | S | *Plexaura kuna* |

| | | |
|---|---|---|
| AY126414 | S | *Plexaurella dichotoma* |
| AY683084 | S | *Plexaurella fusifera* |
| AY126413 | S | *Plexaurella grisea* |
| AY126415 | S | *Plexaurella nutans* |
| KC788276 | D | *Plumarella* sp. |
| HG917043 | S | *Psammogorgia* cf. *arbuscula* |
| AY126401 | S | *Pseudoplexaura crucis* |
| AY683062 | S | *Pseudoplexaura porosa* |
| GQ342524 | S | *Pseudoplexaura wagenaari* |
| AY126421 | S | *Pseudopterogorgia acerosa* |
| AY126423 | S | *Pseudopterogorgia americana* |
| AY126422 | S | *Pseudopterogorgia elisabethae* |
| AY126403 | S | *Pterogorgia anceps* |
| AY126402 | S | *Pterogorgia citrina* |
| KC984590 | M | *Scleracis guadalupensis* |
| KC984579 | D | *Scleraxonia* sp. |
| KC984605 | D | *Sibogagorgia cauliflora* |
| KC984585 | D | *Stolonifera* sp1 |
| KC984591 | D | *Swiftia casta* |
| KC984582 | M | *Swiftia exserta* |
| KC984596 | P | *Swiftia koreni* |
| KC984598 | D | *Swiftia pallida* |
